# Fungal:bacterial biomass balance links environmental gradients to soil respiration across a forest-to-marsh transition

**DOI:** 10.64898/2026.09.23.753783

**Authors:** Courtney E. Mathers, Lillian R. Aoki, Brendan J. M. Bohannan, Josh Roering, Jacob Searcy, Lucas C. R. Silva

## Abstract

Soil microbes regulate whether carbon is retained in soils or returned to the atmosphere through respiration, but the extent to which microbial community characteristics improve the prediction of heterotrophic soil CO_2_ production beyond predictions by environmental controls remains unclear. We tested this across a topographically structured forest-to-marsh gradient in coastal Oregon by measuring heterotrophic soil respiration, soil physicochemical properties, PLFA-based microbial biomass, metagenomic taxonomic composition, and functional gene-based trait indicators. Across the gradient, soil moisture increased from forest to marsh, while mineral soil and organic matter C:N decreased. These environmental shifts were accompanied by strong but uneven microbial responses: total microbial, fungal, and bacterial biomass declined from forest to marsh, taxonomic composition showed the strongest structuring by environmental conditions, and functional gene-based indicators showed mixed relationships with the gradient. Environmental model comparisons identified soil moisture and organic layer C as the strongest baseline predictors of respiration. Among microbial descriptors, only a small subset improved respiration prediction beyond this environmental baseline. The fungal:bacterial (F:B) biomass ratio produced the largest increase in model fit and the greatest reduction in AICc, whereas the best taxonomic and functional-gene predictors yielded more minor gains. Our results showed that the microbial descriptors most responsive to environmental gradients were not the ones most useful for predicting soil respiration; instead, a relatively simple biomass-partitioning metric captured respiration-relevant microbial variation more effectively than finer taxonomic and genomic descriptors. This suggests that F:B ratio may be especially useful for representing respiration-relevant microbial variation in local landscape-scale studies and for future carbon cycle models applied across heterogeneous ecosystem transition zones.

## Introduction

Soil organic carbon (SOC) represents the largest terrestrial reservoir of carbon (C), and its fate is closely governed by the metabolic activity of soil microbes (Crowther et al., 2019). By decomposing soil organic matter (SOM), microbes regulate whether carbon is retained in soils through biomass formation and microbial byproducts or returned to the atmosphere as CO_2_ through respiration. Because soil CO_2_ respiration is one of the largest carbon fluxes between the biosphere and atmosphere, even modest changes in microbial activity can have consequences for ecosystem carbon balance and climate feedbacks (Bond-Lamberty et al., 2018; Friedlingstein et al., 2022). Understanding the drivers of soil CO_2_ respiration is therefore essential for predicting whether soils function as carbon sources or sinks. Current Earth System Models (ESMs) of soil C cycling rely primarily on environmental variables related to climate, vegetation, and soil physicochemical characteristics, and the incorporation of variables representing the microbial community has been highlighted as a priority for improving ESM predictions (Lennon et al., 2024; Wieder et al., 2015a). Despite the central role of microbes in regulating soil respiration, we still lack a clear understanding of which microbial community characteristics are most useful for explaining variation in soil respiration rates and whether microbial information consistently adds explanatory power beyond environmental variables.

Much of our mechanistic understanding of microbial metabolism comes from laboratory and incubation studies showing how soil moisture, temperature, pH, and substrate availability constrain microbial activity (Schimel and Schaeffer, 2012; Moyano et al., 2013). These physicochemical factors regulate microbial metabolism by altering enzyme activity, oxygen availability, nutrient diffusion, and the quantity and accessibility of substrates available for microbial uptake and growth (Burns et al., 2013; Moyano et al., 2013). However, relationships observed under controlled conditions do not scale directly to ecosystem respiration, because environmental variables alone do not capture how microbial communities respond to variation in local soil conditions or how resulting shifts in community structure influence carbon processing and retention in soils (Wieder et al., 2015b; Chandel et al., 2023). Such shifts can alter carbon fate by changing decomposition pathways, growth efficiency, and the balance between carbon loss through respiration and carbon retention in biomass and necromass (Sokol et al., 2022; Camenzind et al., 2023). Yet because soil respiration is an aggregate process emerging from the activity of many microbial populations and functional groups, not all dimensions of microbial community structure are equally likely to improve its prediction (Graham et al., 2016; Hanson et al., 2000). A central challenge, therefore, is to identify which microbial descriptors explain variation in soil respiration beyond what is already captured by environmental variables. Here, we focus on three broad classes of microbial descriptors: biomass-based, taxonomic, and functional gene-based trait indicators.

Biomass-based indicators provide one relatively simple way to represent microbial community structure in relation to soil carbon cycling. Total microbial biomass has been used as a community-scale measure because, as the living microbial carbon pool, it reflects the extent to which assimilated carbon is retained in biomass rather than immediately returned to the atmosphere as CO_2_ (Salazar-Villegas et al., 2016). However, total biomass alone does not distinguish among microbial groups that differ in growth rate, decomposition strategy, and their contributions to carbon retention, which may help explain why biomass-respiration relationships are often context-dependent (Wang et al. 2003; Rui et al., 2016; Yang et al., 2019). Another common biomass-based metric is the fungal:bacterial (F:B) ratio, which reflects how microbial biomass is partitioned between two major decomposer groups often associated with contrasting growth rates, substrate use, and residue formation pathways, with fungi generally considered slower growing than bacteria (Jastrow et al. 2007; Strickland and Rousk, 2010). Higher F:B ratios have been linked to altered carbon cycling, including lower litter-derived CO_2_ loss and greater incorporation of litter-derived carbon into soil organic matter, consistent with greater carbon storage potential in more fungal-dominated communities (Malik et al., 2016). Together, these biomass-based metrics provide tractable community-scale indicators of microbial carbon allocation and decomposer dominance, but their coarse resolution may limit how clearly they distinguish the mechanisms linking microbial communities to soil respiration.

Taxonomic approaches provide another way to represent microbial community structure in relation to soil carbon cycling. In some cases, taxonomic composition has explained unique variation in soil respiration beyond soil properties and broader microbial attributes such as biomass or richness, suggesting that community composition can capture ecologically meaningful differences in microbial carbon-processing strategies (Liu et al., 2018, Liu et al., 2020). In bacterial communities, one common contrast is between taxa associated with more copiotrophic versus oligotrophic strategies, a distinction reflecting differences in whether carbon is processed through faster, opportunistic use of readily available substrates or slower turnover under more resource-limited conditions (Fierer et al., 2007; Ho et al., 2017). In fungal communities, variation in the relative dominance of ectomycorrhizal (EM) versus arbuscular mycorrhizal (AM) associations has likewise been linked to contrasting nutrient economies and soil carbon storage (Averill et al., 2014; Averill and Hawkes, 2016). Taxonomic groupings may therefore provide ecological context for interpreting variation in soil respiration, while remaining indirect indicators of the mechanisms through which microbes influence carbon fate.

Trait-based approaches provide a third way to represent microbial community structure in relation to soil carbon cycling by focusing on functional traits that influence how microbes acquire resources, tolerate stress, grow, and contribute to ecosystem processes (Violle et al., 2007; Krause et al., 2014). Metagenomic data provide one way to infer such traits from community gene content, using genes that code for proteins that control biochemical pathways as indicators of the traits and capabilities represented within microbial communities (Fierer et al., 2012). One example is the Y-A-S framework proposed by Malik et al. (2020), which has been used to interpret microbial life-history strategies in relation to soil carbon cycling through tradeoffs among growth yield (Y), resource acquisition (A), and stress tolerance (S) (Broderick et al., 2025; Jones et al., 2025; Nelson et al., 2024). Another is the abundance of genes encoding carbohydrate-active enzymes (CAZymes), which can be used to characterize the community’s potential to decompose organic matter derived from plants, fungi, and bacteria and to interpret decomposition-related soil carbon cycling across environmental gradients (Huang et al., 2024; Xiong et al., 2023). Because these gene-based indicators are tied to carbon allocation and decomposition potential, they may offer a more mechanistic framework for linking microbial communities to soil respiration than biomass or taxonomy. However, gene abundance reflects functional potential rather than realized activity, because expression and process rates remain constrained by environmental context and substrate accessibility (Blagodatskaya et al., 2021; Rocca et al., 2015). Gene-based functional indicators may thus provide added functional resolution, but whether they improve prediction of soil respiration beyond simpler microbial community-scale descriptors remains an open question (Liu et al., 2020; Malik et al., 2020).

To address these questions, we measured soil respiration rates across a topographically structured forest-to-marsh gradient in coastal Oregon, where soil saturation, organic matter accumulation, and microbial community composition varies with landscape position. Environmental gradients of this kind are useful for testing microbial controls on respiration because they generate correlated shifts in abiotic conditions, vegetation inputs, and microbial community attributes across space, allowing us to ask whether microbial descriptors improve prediction beyond the environmental structure they track. Within this gradient, the ecotone marks the zone where upland and marsh influences overlap, creating especially strong local heterogeneity in hydrology, organic matter inputs, and soil conditions over short spatial scales. We evaluated whether three classes of microbial descriptors, biomass-based, taxonomic, and gene-based trait indicators, improved prediction of soil respiration beyond environmental variables alone. We hypothesized that 1) environmental conditions and microbial community structure would vary systematically across the gradient, with hydrology and vegetation-related differences in organic matter expected to be important drivers of that variation, and 2) microbial community characteristics would explain additional variation in soil respiration beyond what was captured by environmental variables alone. We then compared microbial descriptor classes to test whether finer taxonomic or gene-based trait indicators provided additional explanatory power beyond simpler biomass-based measures.

## Methods

### Study site and sample collection

In winter 2023, we collected soil samples along five transects extending from a forested upland hillslope into Kunz Marsh, a tidal high marsh, within the South Slough National Estuarine Research Reserve on the coast of Oregon, USA. Each transect descended from approximately 40 m elevation in the upland forest to near sea level in the marsh and spanned strong contrasts in topography, vegetation, and hydrology that were expected to influence drainage and organic matter accumulation. Because marsh soil conditions vary with daily and seasonal tides, sampling was conducted during a low-tide period to improve access to high-marsh soils under relatively drained conditions. Five sampling points were established approximately 15 m apart along each transect, with two points in the forest, one in the ecotone, and two in the high marsh (Figure 1). Forest and ecotone sampling points were dominated by EM-associated trees with understory ferns and ericoid shrubs, whereas high-marsh sampling points were sedge-dominated. Soils at our study site consisted of Inceptisols in the steep upland forest slopes, a mosaic of Inceptisols and Histosols occurring along the ecotone, and Histosols and Entisols in the marsh. At each sampling point, we collected and composited three mineral-soil samples (0-20 cm depth) using a soil auger, resulting in a total of 25 composite soil samples, with five transect replicates per landscape position. We also collected organic layer material consisting of surface plant litter above the mineral soil at each sampling point to capture landscape-position differences in aboveground organic matter inputs across the gradient. All samples were stored in a cooler in the field and transported to the laboratory the same day.

**Figure 1:**
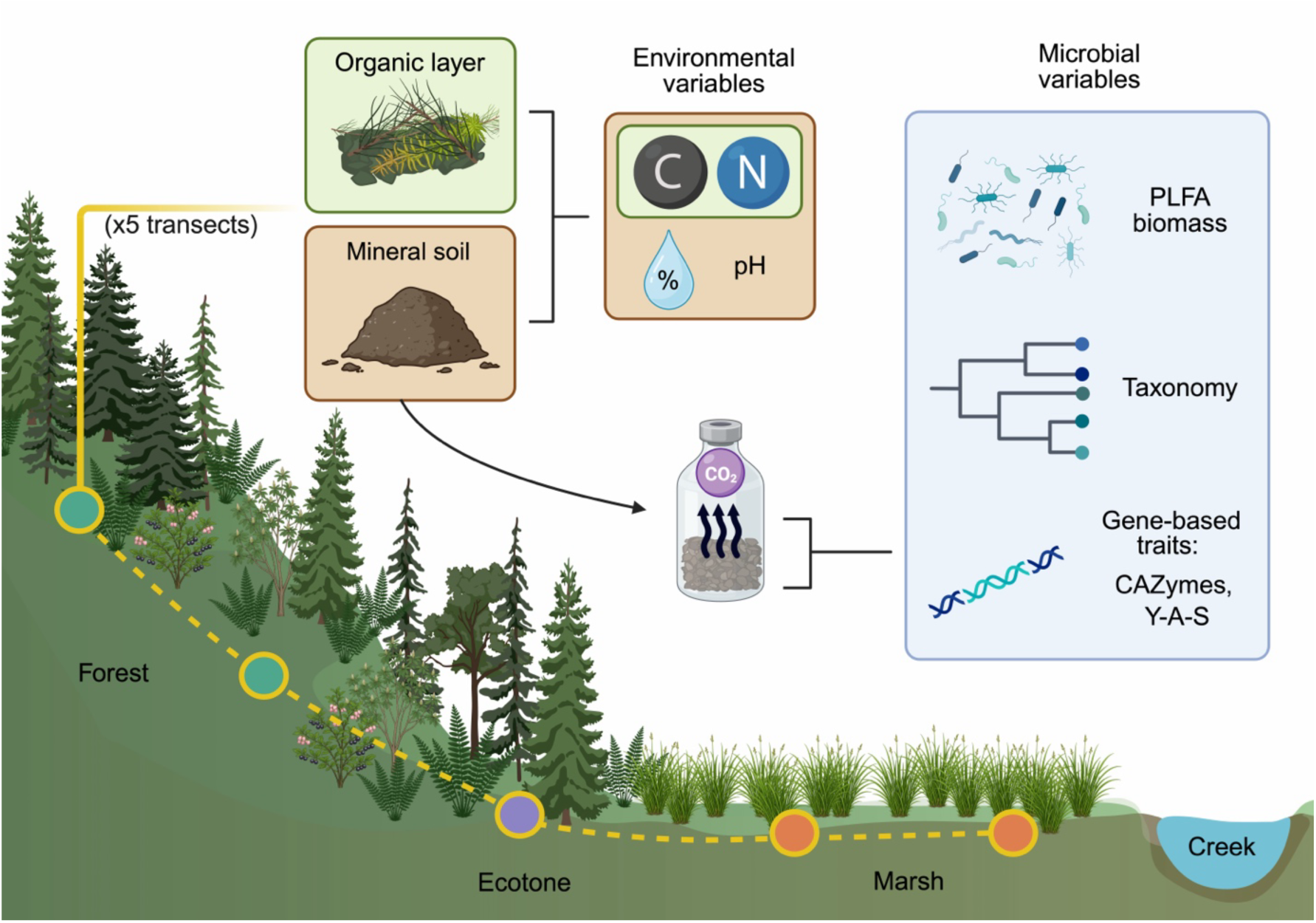
Study site cross section. Environmental variables included measures of soil moisture, soil pH, and %C and %N for both mineral soil and organic layer samples. Microbial variables included measures of community biomasses (PLFA), metagenomic taxonomy, CAZymes, and Y-A-S traits. Created with BioRender.com.

### Soil respiration measurements

Upon returning to the lab, approximately 15 g of field moist homogenized samples were placed in 120 mL glass incubation vials after removing and large roots and plant debris. Because root material was removed prior to incubation, resulting CO_2_ fluxes reflect heterotrophic, microbially mediated decomposition of SOM rather than total field soil respiration. Samples were incubated at field moisture to preserve differences in sample hydrologic conditions and aeration. Vials were flushed with CO_2_-free air for 15 minutes, and incubated at 10°C, the average soil temperature of all sampling points at the time of sampling. Headspace CO_2_ and CH_4_ were measured after 1, 3, and 5 h on a gas chromatograph equipped with a flame ionization detector. Gas concentrations were converted to flux rates following Bridgham and Ye (2013), and CO_2_ production rates were calculated from the linear slope of the time series and expressed as μmol CO_2_ produced per g of dry soil per hour. CH_4_ did not increase during incubation, indicating that aerobic conditions were maintained. Percent soil moisture was then calculated on a mass basis after oven drying.

### Soil chemical analyses

Soil samples were separated into three sets of sub-samples. One set was oven-dried, sieved to 2 mm, ground, and used to analyze total SOC and total N with an elemental analyzer. Organic layer samples were also ground and analyzed for total organic C and total N in the same manner. Soil pH was measured for mineral layer samples with a 1:1 soil:deionized water slurry. The second set of soil sub-samples were sent overnight on ice to Ward Laboratories, Inc (Kearney, NE) for phospholipid fatty acid analysis (PLFA) to estimate bacterial and fungal biomass. Two samples were excluded because PLFA measurements were unavailable after laboratory processing. Because PLFA does not capture archaeal biomass, archaeal patterns were evaluated only with metagenomic data.

### Sequencing and quality control

The final set of soil subsamples was stored at -20°C until total DNA was extracted from 0.25 g of each sample using the DNeasy PowerLyzer PowerSoil Kit (Qiagen, Germantown, MD, USA) following the manufacturer’s protocol. Metagenomic sequencing libraries were prepared using the Illumina DNA Prep Kit and sequenced by the University of Oregon Genomics and Cell Characterization Core Facility on an Illumina NovaSeq 6000 with an S4 flow cell (PE 150 bp). Approximately 21–44 Gbp of raw sequence data were generated per sample, with a mean of 31.4 Gbp per sample. Read quality was initially assessed with FastQC (v0.12.1), and one ecotone sample was excluded from downstream analyses because of low sequencing yield (0.12 Gbp). Reads were then quality trimmed and adapter sequences removed with TrimGalore (v0.6.1). After trimming, 21–42 Gbp of high-quality sequence data remained per sample.

### Assembly and taxonomic classification

For each sample, trimmed reads were assembled into contigs using the de novo de Bruijn assembler MEGAHIT (v1.2.9; minimum k-mer 27, maximum k-mer 127, step size 10), and assembly quality was assessed with QUAST (v5.3.0). Trimmed reads were subsequently mapped back to assemblies with Bowtie2 (v2.5.4), and the resulting alignments were processed into sorted and indexed BAM files with samtools (v1.23) for downstream coverage profiling and gene-level abundance estimation. We taxonomically classified trimmed paired-end reads with Kraken2 (v2.1.2), which assigns reads to taxonomic nodes based on exact k-mer matches (Wood et al., 2019). The Kraken2 Standard and PlusPF database was downloaded in January 2026 to capture bacterial, archaeal, viral, and fungal sequence diversity. Bracken (v2.8) was then used to re-estimate relative abundances at class level from Kraken2 output, correcting for biases associated with read length and genome size (Lu et al., 2017).

### Gene prediction and functional trait annotation

We performed gene prediction in Anvi’o (v9) using Prodigal (v2.6.3) in metagenomic mode on assembled contigs >1000 bp. Gene-level coverage was then estimated in Anvi’o from the mapped-read alignments described above, and these coverage values served as the basis for all downstream gene-based trait abundance calculations. In this study, gene coverage was treated as an index of function abundance because it reflects the amount of mapped metagenomic signal associated with each annotated gene or gene group, allowing community-level functional potential to be compared among samples.

We annotated predicted genes with Kyoto Encyclopedia of Genes and Genomes (KEGG) Orthologs using Anvi’o, which assigns KEGG Ortholog (KO) annotations through HMM-based searches against the KOfam database. Within each sample, genes assigned to the same KO were aggregated to generate KO-level abundance estimates. KOs were then linked to MicroTrait’s Y-A-S framework using a curated KO-to-trait mapping table from Broderick et al. (2025), which was adapted from Karaoz and Brodie’s (2022) original MicroTrait framework for use with assembled metagenomic contigs rather than metagenome-assembled genomes. Annotated KOs were assigned to either Resource Use/Yield (Y), Resource Acquisition (A), or Stress Tolerance (S) categories, which consisted of 88, 864, and 132 unique KO ids, respectively. KO-level abundances were subsequently summed within each trait category to estimate Y-A-S trait abundances for each sample.

We also annotated predicted genes for carbohydrate-active enzymes (CAZymes) using dbCAN3 (Zheng et al. 2023). Genes assigned to the same CAZyme family were aggregated to estimate family-level abundance, and annotated families were grouped into broader substrate-degradation categories following López-Mondéjar et al. (2020), based on functional assignments in the CAZy database (http://www.cazy.org); full family-to-group assignments are provided in Appendix S1:Table 1. These categories included cellulose, hemicellulose, and lignin associated with plant biomass, chitin and glucans associated with fungal biomass, and peptidoglycan associated with bacterial biomass. This grouping framework allowed CAZyme annotations to be interpreted in terms of broader substrate targets and related microbial functional potential to ecologically relevant sources of SOM (Huang et al. 2024, Xiong et al. 2023, Zuo et al. 2026). To account for differences in sequencing depth among samples, summed gene coverage values for KO-derived trait categories and CAZyme groups were normalized by total mapped reads and expressed as counts per million mapped reads (CPM).

**Table 1:** Comparison of the strongest biomass-based, taxonomic, and gene-based microbial add-ons across environmental baseline models explaining soil heterotrophic respiration.

| Model | Predictors | n | $R^2_{\text{adj}}$ | $\Delta\text{AICc}$ |
| --- | --- | --- | --- | --- |
| **Best environmental base model** | Soil moisture + organic layer C | 25 | 0.653 |  |
| Best biomass add-on | Envir. base + F:B | 23 | 0.692 | -1.764 |
| Best taxonomic add-on | Envir. base + Methanomicrobia | 24 | 0.687 | -0.601 |
| Best gene-based trait add-on | Envir. base + hemicellulose | 24 | 0.658 | 1.562 |
| **Soil C base model** | Soil moisture + soil C (log) | 25 | 0.461 |  |
| Best biomass add-on | Soil C base + F:B | 23 | 0.509 | -3.028 |
| Best taxonomic add-on | Soil C base + fungi PCoA | 24 | 0.500 | -0.144 |
| Best gene-based trait add-on | Soil C base + hemicellulose | 24 | 0.508 | -0.520 |
| **PCoA environmental base model** | Soil moisture + Envir. PCoA1 | 25 | 0.515 |  |
| Best biomass add-on | PCoA base + F:B | 23 | 0.522 | -1.716 |
| Best taxonomic add-on | PCoA base + fungi PCoA | 24 | 0.575 | -1.429 |
| Best gene-based trait add-on | PCoA base + peptidoglycan | 24 | 0.541 | 0.438 |

## Statistical analyses

All analyses were conducted in R (v4.5.2). Variables were log-transformed where necessary prior to statistical testing and model fitting, and statistical significance was evaluated at α = 0.05. Unless otherwise indicated, we show raw values in figures for interpretability, while transformed values were used for formal tests where appropriate.

We organized microbial variables into three descriptor classes: biomass-related variables from PLFA analyses, taxonomic variables derived from relative abundance tables of the 12 most abundant bacterial, fungal, and archaeal classes in each sample, and gene-related trait variables, including Y-A-S trait abundances and grouped CAZyme abundances. Separate Bray-Curtis distance matrices were constructed from bacterial, fungal, and archaeal relative-abundance tables using the vegan package (v2.7.2). Principal coordinates analysis (PCoA) was used to summarize taxonomic variation, community differences among landscape positions were evaluated with permutational multivariate analysis of variance (PERMANOVA), and the first axis from each PCoA was retained for downstream correlation analyses and models.

Several bacterial, fungal, and archaeal classes were also selected as candidate variables in correlation analyses and respiration models based on their prevalence across samples and their ecological relevance to soil C and N cycling, thereby prioritizing biologically interpretable variables over coincidental statistical relationships. Alphaproteobacteria were selected for their oligotrophic traits, while Gammaproteobacteria were chosen for their copiotrophy (Kurm et al., 2017). Actinomycetes were also for their roles in nitrogen fixation and the breakdown of recalcitrant SOM inputs such as lignin, cellulose, and chitin (Bhatti et al., 2017). Selected fungal classes included Glomeromycetes, which comprise AM fungi, and Agaricomycetes, which includes many EM taxa and numerous other mushroom-forming saprotrophs (Tedersoo et al., 2010). Archaeal classes included Methanomicrobia and Nitrososphaeria for their roles in methanogenesis (Hofmann et al., 2016) and ammonia-oxidation (Saghaï et al., 2022).

We evaluated differences in soil respiration rates, environmental variables, and microbial-related variables among forest, ecotone, and marsh landscape positions using analysis of variance (ANOVA) followed by Tukey HSD tests for pairwise comparisons. Relationships among variables were evaluated using linear regression, and the strength and direction of bivariate relationships were quantified using Pearson correlation coefficients (*r*). Because sample size was limited (n = 25), we restricted model comparisons to low-complexity linear models to reduce overfitting. We selected candidate environmental variables to represent major dimensions of the topographically structured gradient, including percent soil moisture, soil pH, mineral soil organic %C, mineral soil C:N, organic layer %C, and organic layer C:N. For each microbial response variable, all possible one- and two-variable combinations of these environmental predictors were fit using linear regression and ranked using R²_adj_ and corrected Akaike Information Criterion (AICc), and the model with the lowest AICc was retained as the best-supported model. To address our first hypothesis, we then grouped best-supported models into biomass-related, taxonomic, and gene-based functional trait categories to compare the strength of environmental structuring across different dimensions of the microbial community.

We then used the same candidate-model approach to evaluate predictors of soil respiration. A baseline environmental model for respiration was first identified as the best-supported environmental-only model. Two additional environmental baselines, one based on soil C and one based on environmental PCoA1 derived from soil pH, soil C, soil C:N, organic layer C, and organic layer C:N, were retained as sensitivity checks to assess whether microbial model improvements were robust to baseline model specification. Microbial variables were added individually to base models to test whether they explained additional variation in respiration beyond the environmental baselines. We used changes in R²_adj_ and AICc to quantify the extent to which each microbial variable improved model fit relative to the environmental baseline.

## Results

### Soil physiochemical characteristics and soil respiration were related to landscape position

Measured soil physicochemical properties varied across landscape positions, with soil moisture increasing from forest down to marsh (ANOVA, *p* = 0.002) and both organic layer C:N and mineral soil C:N decreasing along the gradient (*p* = 0.001 and *p* < 0.001) (see Appendix S1: Figure 1 for all environmental variables across landscape positions). Soil pH ranged from approximately 3.7 to 5.9 and varied across landscape positions (*p* = 0.001), with the lowest mean pH in ecotone soils (4.04 ± 0.19, mean ± SE) and higher values in forest (4.52 ± 0.08) and marsh soils (4.96 ± 0.16). Respiration rates ranged from 0.025 to 0.134 µmol CO_2_ g^-1^ soil h^-1^ and were generally lower in the forest and higher in the ecotone; however no significant differences in respiration were identified across landscape positions due to high within-group variation (*p* = 0.823; Figure 2A). Soil respiration was most positively correlated with soil C (*r* = 0.70, *p* < 0.001) and soil moisture (*r* = 0.59, *p* < 0.01), whereas soil pH, organic layer C, and organic layer C:N showed weaker relationships with respiration (Figure 2B).

**Figure 2:**
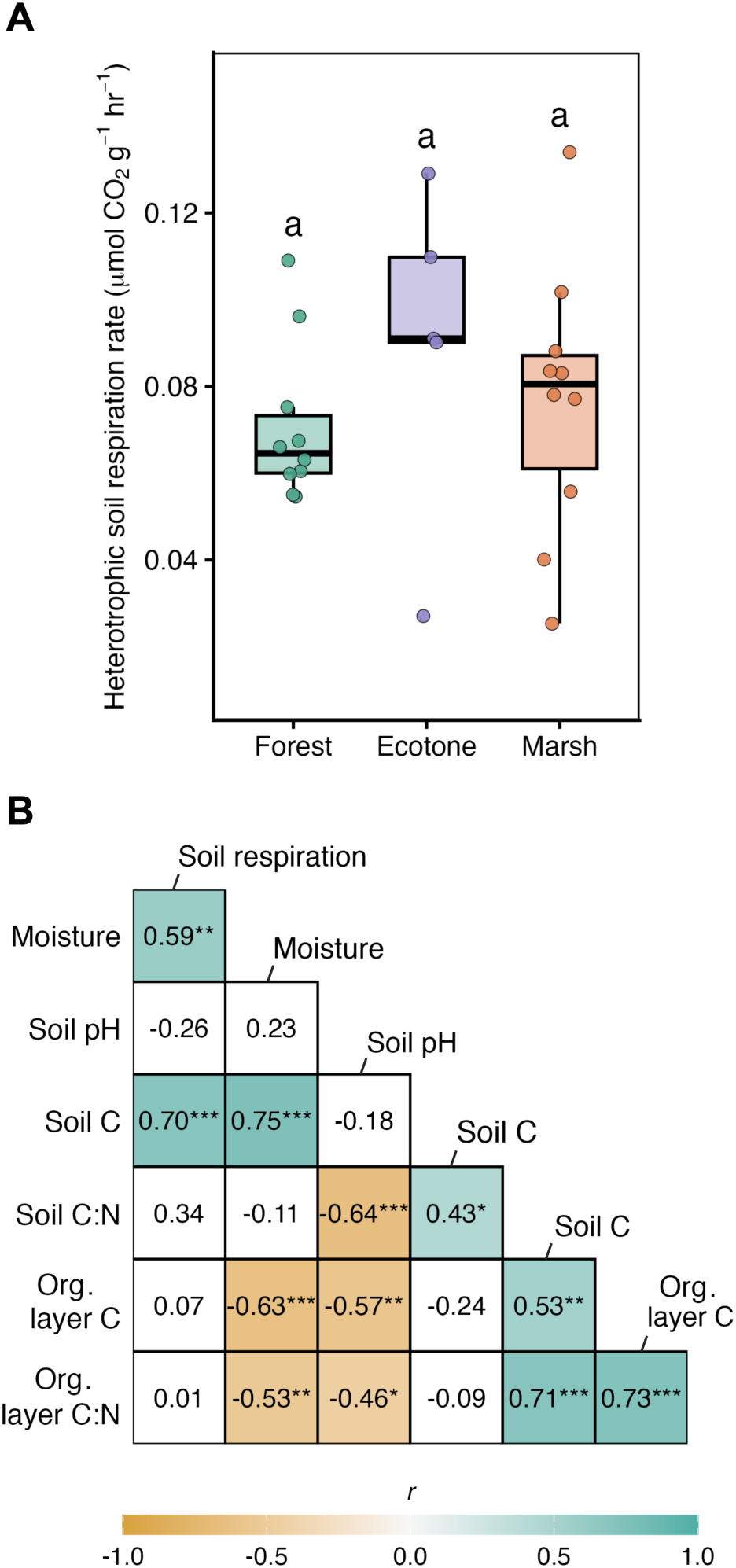
(A) Soil respiration rates across landscape positions. Letters denote Tukey HSD groupings from pairwise comparisons. (B) Pearson correlation matrix for soil respiration rates and environmental variables. Values in each cell are Pearson correlation coefficients (*r*), with asterisks denoting significance (* *p* < 0.05, ** *p* < 0.01, *** *p* < 0.001).

### Microbial community characteristics were related to landscape position

Landscape position was statistically associated with multiple attributes of the microbial community, but the strength of that relationship differed among biomass-based, taxonomic, and gene-based variables. Total microbial biomass, total fungal biomass, and total bacterial biomass were all highest in the forest and declined through the ecotone into the marsh (*p* <= 0.004). Although we expected fungal dominance to be greatest in the forest, F:B ratios did not differ among landscape positions (*p* = 0.679) and exhibited substantial within-group variation, particularly in ecotone samples, which included both the lowest and highest F:B values observed across the site (Appendix S1:Figure 2).

Taxonomic microbial community composition from metagenomic read classification also varied across the forest-to-marsh gradient. PCoA of Bray-Curtis dissimilarities showed clear shifts in bacterial, fungal, and archaeal community composition across landscape positions (Appendix S1:Figures 3-5). Bacterial and archaeal community composition differed between marsh and both upland positions (PERMANOVA, *p* < 0.03), whereas forest and ecotone communities did not. Fungal communities showed weaker pairwise separation, with only the forest-marsh comparison significant (*p* = 0.001), indicating that the ecotone was compositionally intermediate between the two end-member habitats (Appendix S1: Figure 6). All landscape positions shared the same four most abundant bacterial classes, with Alphaproteobacteria, Actinomycetes, Gammaproteobacteria, and Betaproteobacteria dominating across samples. In contrast, fungal and archaeal classes shifted more clearly across the gradient. Glomeromycetes increased from forest to marsh soils (*p* < 0.04; Appendix S1: Figure 7), whereas Agaricomycetes were more abundant in forest and ecotone positions (*p* < 0.001), consistent with the expected shift from EM- to AM-dominated plant communities. Archaeal reads were also relatively more abundant in marsh soils, including higher abundance of Methanomicrobia (*p* = 0.001), although several ecotone samples and one forest sample showed elevated archaeal abundance as well. Together, these results indicate that landscape position influenced not only broad multivariate composition, but also the abundance of specific microbial groups with distinct ecological roles in soil carbon cycling.

**Figure 3:**
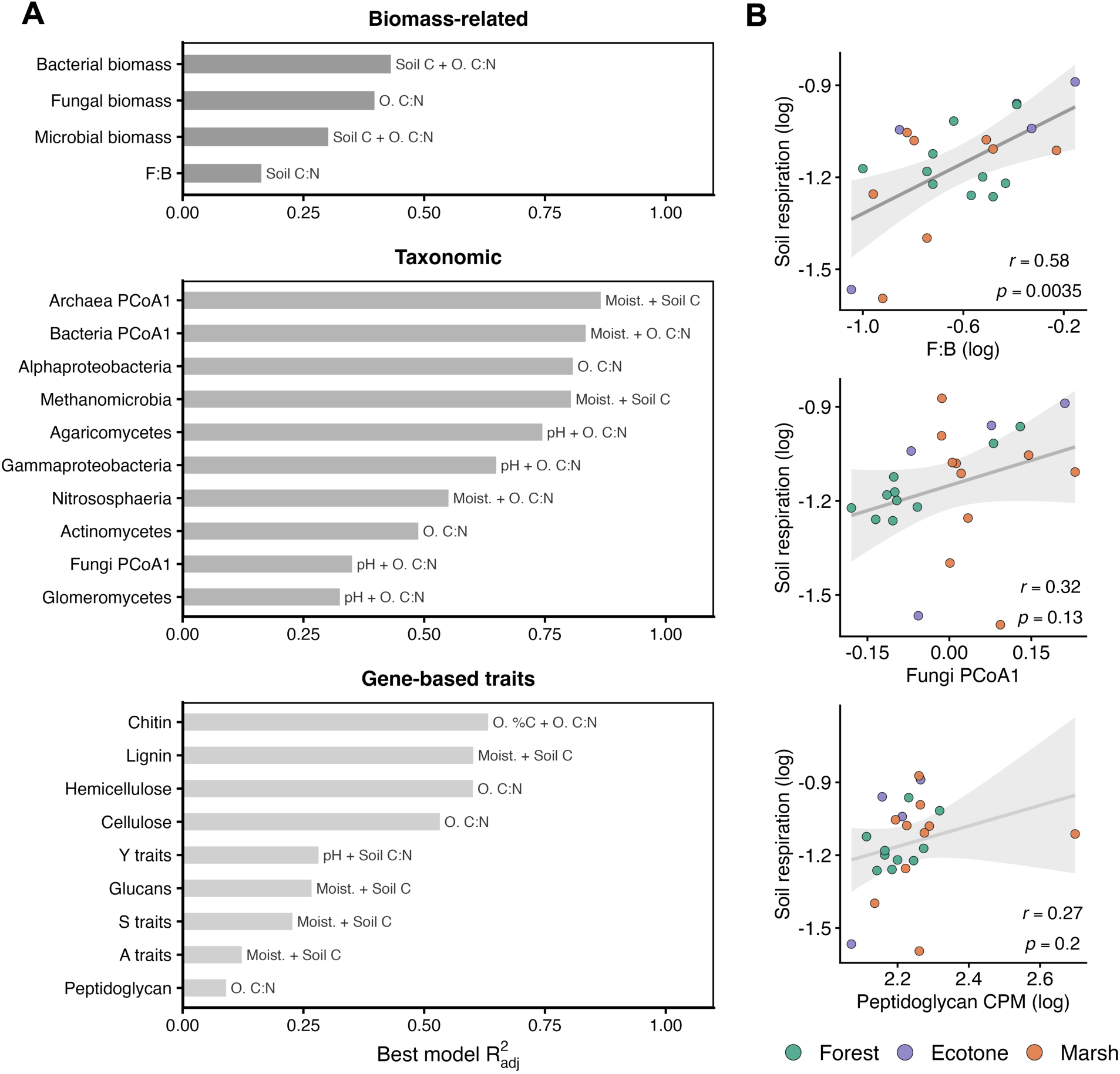
Environmental predictors of microbial response variables and strongest microbial associations with soil heterotrophic respiration. (A) For each biomass-related, taxonomic, and gene-based trait response variable, bar length shows R^2^_adj_ of the environmental model with the lowest AICc among the candidate one- and two-predictor models. Text labels identify the environmental predictor(s) retained in the best-supported model. Moisture and organic layer are abbreviated as “Moist.” And “O.”, respectively. (B) Microbial predictors showing the strongest associations with log-transformed soil respiration (original units: μmol CO_2_ g^-1^ hr^-1^) within each of the three categories. Lines show fitted linear relationships with 95% confidence intervals, and points are colored by landscape position.

Gene-based trait profiles showed much weaker landscape structuring than taxonomic composition. Y and S trait abundances did not differ significantly among forest, ecotone, and marsh soils, and although A traits showed a modest overall effect of landscape position, no individual Tukey HSD pairwise comparisons were significant (Appendix S1:Figure 8). These results suggest that broad Y-A-S functional strategies were comparatively conserved across the gradient, even as biomass and community taxonomic composition shifted substantially among positions. Landscape position was associated with grouped CAZyme abundances more strongly than Y-A-S traits (Appendix S1:Figures 9 and 10). Cellulose, hemicellulose, lignin, glucans, and chitin all varied significantly among landscape positions (*p* ≤ 0.03), whereas peptidoglycan did not (*p* = 0.188). Overall, plant-associated CAZyme groups shifted from more lignin-rich forest profiles toward more cellulose- and hemicellulose-rich marsh profiles, while fungal biomass-associated glucans and chitin showed contrasting patterns and peptidoglycan remained comparatively constant. These results indicate strong landscape-related shifts in CAZyme groups associated with plant and fungal biomass despite weak structuring of broad Y-A-S categories.

### Environmental variables explained variation in microbial community structure

To identify which environmental variables were most strongly associated with microbial community structure, we compared best-supported environmental models for each biomass-based, taxonomic, and gene-based response variable (Figure 3A). Taxonomic responses were the most strongly structured by environmental variation, with the highest mean best-model R²_adj_ (0.64), substantially exceeding biomass-related (0.32) and gene-based (0.37) mean responses. The strongest individual taxonomic fits were observed for Archaea PCoA1 (R²_adj_ = 0.86), Bacteria PCoA1 (0.83), and Alphaproteobacteria (0.81), and their best-supported models repeatedly included moisture together with organic layer or mineral-soil stoichiometric variables. Correlation patterns were consistent with these results, indicating that moisture and especially organic layer C:N were the most common environmental correlates of taxonomic composition (Appendix S1: Figure 11). Biomass-related variables showed more moderate environmental structure. Best-model comparisons indicated that total biomass measures were best explained by combinations of soil C and organic layer C:N, and F:B was best explained by mineral soil C:N (Figure 3A). Correlations showed the same general pattern, with biomass measures most strongly associated with soil stoichiometry while showing weaker associations with soil moisture than taxonomic variables (Appendix S1: Figure 11).

Gene-based trait estimates were more variable in their environmental patterning. Broad Y-A-S traits showed weak to moderate structure, with best-model R²_adj_ values ranging from 0.12 for A traits to 0.28 for Y traits. In contrast, grouped CAZyme abundances showed clearer environmental patterning, particularly for chitin and plant-associated groups. Cellulose, hemicellulose, and lignin CAZyme abundances were all moderately well explained by environmental models (R²_adj_ = 0.53–0.60), with correlation patterns indicating that cellulose and hemicellulose were associated with wetter, lower-C:N conditions, whereas lignin CAymes were associated with drier, higher-C:N conditions (Appendix S1: Figure 11). Chitin showed moderate environmental structure, while peptidoglycan and glucans were comparatively weak.

Although taxonomic variables were more strongly structured by environmental conditions than biomass-related variables, none showed a significant relationship with soil respiration. Methanomicrobia and fungi PCoA1 had the strongest taxonomic respiration correlations (*r* = 0.34 and 0.32), and peptidoglycan had the strongest correlation among gene-based traits (*r* = 0.27), but all these relationships were weak and not significant. By contrast, the fungal:bacterial biomass ratio was the only microbial variable significantly associated with soil respiration (*r* = 0.58, *p* < 0.01; Figure 3B).

Together, these results support our first hypothesis that environmental conditions and microbial community structure varied systematically across the forest-to-marsh gradient. Soil moisture and especially organic layer C:N were the most consistent environmental correlates of microbial community characteristics, but the strength of this structuring was greatest for taxonomic composition, weaker for biomass-based metrics, and more variable across gene-based trait categories.

### Soil respiration model testing

Environmental baseline model comparisons showed that soil respiration across the forest-to-marsh gradient was best explained by a baseline environmental model including soil moisture and organic layer C (R²_adj_ = 0.65, AICc = -34.12; Table 1). Adding landscape position did not significantly improve model fit (*p* = 0.85), indicating that respiration differences among forest, ecotone, and marsh soils were explained primarily by measured environmental variation rather than by landscape category itself. To assess sensitivity to baseline model specification, we also retained two alternative environmental baselines for microbial add-on comparisons: one based on soil C, and one based on environmental PCoA1 derived from soil pH, soil C, soil C:N, organic layer C, and organic layer C:N (Table 1).

We evaluated whether microbial variables improved prediction beyond these three environmental baselines by adding individual biomass, taxonomic, and gene-based trait predictors to the baseline models. Among all candidates, F:B biomass ratio consistently produced the strongest improvement across all three baselines (Table 1). By contrast, the best taxonomic add-ons, Methanomicrobia and fungi PCoA1, were more sensitive to baseline specification, with fungi PCoA appearing as the strongest taxonomic add-on in the soil C and environmental PCoA base models. Similarly, the best performing gene-based trait add-ons varied across base models and included hemicellulose and peptidoglycan CAZyme abundances. Taxonomic and gene-based trait add-ons generally produced smaller gains than F:B or failed to improve AICc. The best overall combined model therefore included moisture, organic layer C, and F:B biomass ratio with F:B increasing model fit to R²_adj_ = 0.692 and reducing AICc by -1.764 (Table 1). This pattern was also consistent with bivariate relationships between soil respiration and microbial predictors (Appendix S1:Figure 11), in which F:B biomass ratio and fungi PCoA showed among the strongest positive associations with respiration (*r* = 0.58 and 0.32, respectively; Figure 3B).

Across all microbial add-on models, only a small subset improved model fit relative to the best environmental baseline, with F:B showing the largest gain in explanatory power and model support (Figure 4). Methanomicrobia also improved model support, whereas most other taxonomic and functional-gene predictors either increased AICc or produced only minor gains in R²_adj_ (Appendix S1: Table 2). Follow-up interaction models showed that the effect of F:B did not vary significantly with moisture, organic layer C, organic layer C:N, soil C, soil C:N, or pH (*p* ≥ 0.176 in all cases), indicating that its contribution to respiration was additive rather than strongly contingent on the measured environmental gradients. Thus, our second hypothesis was only partially supported: microbial community characteristics added explanatory power beyond the environmental baseline, but this improvement was driven primarily by F:B biomass ratio and, to a lesser extent, by a small number of taxonomic descriptors.

**Figure 4:**
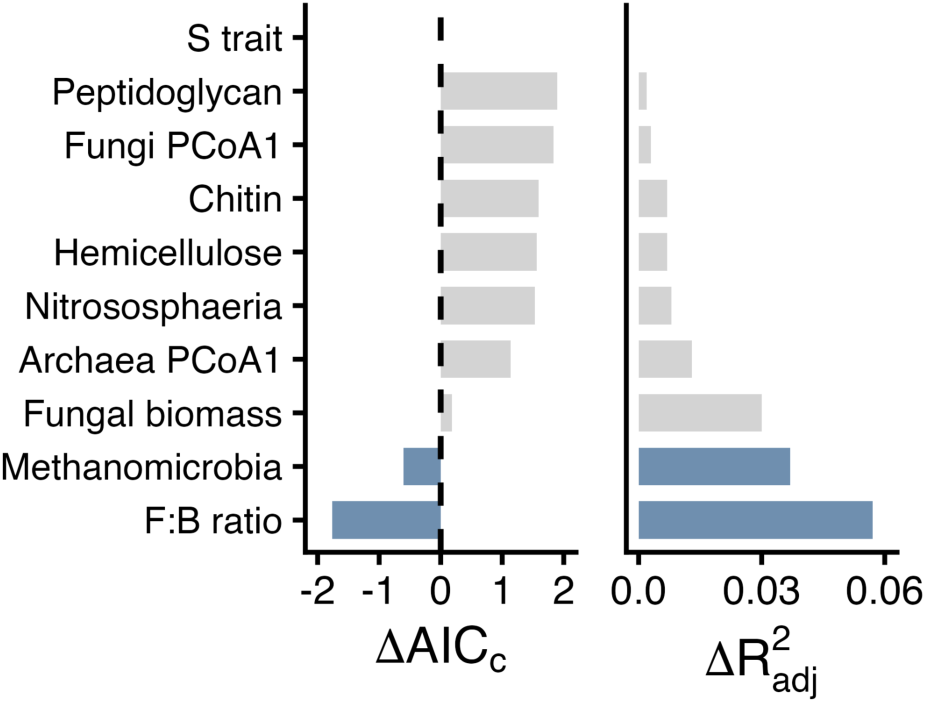
Effects of microbial predictors on soil respiration model performance after addition to the best environmental baseline model (containing soil moisture + organic layer C predictors). Values show changes in AICc and R²_adj_ relative to the corresponding environmental baseline refit on the same subset of samples; only predictors that improved at least one performance metric are shown, and all retained predictors were supported in their log-transformed forms. See Appendix S1: Table 2 for results across all microbial predictors.

## Discussion

We asked whether microbial community descriptors could improve prediction of soil respiration beyond the strong environmental variation imposed by a forest-to-marsh gradient. The best-supported environmental baseline model indicated that respiration was governed primarily by hydrologic conditions and substrate availability, consistent with broader soil-respiration literature emphasizing water availability and plant-derived carbon inputs as key controls on microbial C cycling (Moyano et al., 2013; Schaefer et al., 2009;). Against this strong baseline, F:B biomass ratio was the only microbial descriptor that consistently improved prediction, whereas taxonomic composition and functional gene-based indicators tracked the gradient strongly but added little explanatory power.

The contribution of F:B was especially notable in this regard because it emerged despite only moderate environmental structuring and no clear differences among landscape positions. Its predictive value therefore did not appear to arise from broad categorical differences among forest, ecotone, and marsh soils. Instead, F:B may have integrated finer-scale variation in decomposer balance that was more directly related to respiration than either taxonomic turnover or broad trait-based functional potential. This interpretation is consistent with growing evidence that F:B can capture meaningful variation in soil carbon cycling and improve prediction of soil respiration or related CO_2_ fluxes beyond abiotic drivers alone (Chen et al., 2020; Guasconi et al., 2026). Microbial descriptors are most likely to improve prediction when they capture either rate-limiting steps in decomposition or the fate of carbon after microbial uptake (Schimel and Schaeffer, 2012; Sokol et al., 2022), which may help explain why F:B retained predictive value where finer taxonomic and functional descriptors did not.

The positive association between respiration and F:B indicates that relatively fungal-dominated communities were linked to greater decomposition activity in this system (Figure 3B). This contrasts with the more common association between higher F:B and greater soil C storage or slower turnover (Malik et al., 2016; Six et al., 2006), suggesting that the relationship between F:B and C cycling is strongly context dependent (Bailey et al., 2002; He et al., 2020). A similar positive F:B-respiration relationship has also been reported in managed Scots pine forests, reinforcing that fungal dominance can in some systems be associated with faster C turnover (Guasconi et al., 2026). However, this relationship likely emerged with uneven contributions of different fungal groups. In our data, Glomeromycetes was positively associated with respiration (*r* = 0.23; Appendix S1:Figure 11), whereas Agaricomycetes was (*r* = -0.06) not, indicating that the respiration signal captured by F:B was not driven uniformly across fungal groups. Increases in fungal biomass may also have included saprotrophic fungi with strong extracellular enzymatic capacities, providing a plausible mechanism for the higher respiration observed in more fungal-dominated communities (Burns et al., 2013; Fang et al., 2020).

These uneven fungal contributions likely extended beyond decomposition itself to the persistence and function of the microbial residues in soil C pools. Although F:B is a relative measure, fungal residues may be especially important because fungal necromass often contributes a larger share of total microbial necromass and soil organic carbon than bacterial necromass, including in wetland systems (Li et al., 2015; Wei et al., 2023). EM fungi tend to produce more chemically persistent mycelial residues than AM fungi (Huang et al., 2022), and fungal necromass more broadly contains chemically distinct fast- and slow-cycling pools across taxa (See et al., 2021). Fungal groups may also differ in their contribution to soil carbon retention through aggregate formation and the physical protection of soil organic matter (Rillig et al., 2015; Lehmann et al., 2020). Taken together, these patterns suggest that F:B captured a broader balance between fungal and bacterial influences on soil carbon cycling, integrating processes related to both decomposition and residue persistence.

By comparison, total microbial, fungal, and bacterial biomass tracked the landscape gradient more directly than F:B. Larger standing biomass pools were associated with the drier, forest-influenced end of the transect, where organic matter inputs had higher C:N and were likely more lignin-rich under forest vegetation. However, this stronger environmental patterning did not translate into stronger relationships with soil respiration, suggesting that bulk biomass alone did not capture the aspects of community organization most relevant to heterotrophic CO_2_ production in this system. Consistent with this interpretation, previous studies have shown that microbial biomass and respiration do not necessarily vary together even when respiration responds strongly to substrate conditions (Wang et al., 2003; Rui et al., 2016). Because routine PLFA measurements do not capture archaeal biomass, this limitation may have further weakened the relationship between bulk PLFA biomass and respiration in a system where archaeal abundance increased toward the marsh.

Taxonomic composition responded more strongly to the gradient than biomass measures, with bacterial, fungal, and archaeal communities all shifting significantly and marsh communities consistently the most distinct. Hydrologic and stoichiometric contrasts across the transect reorganized community membership in ways consistent with studies showing that flooding, oxygen limitation, and related soil conditions can strongly restructure microbial composition in wetland-influenced soils (Balasooriya et al., 2008; Peralta et al., 2014). Fungal shifts in taxonomic composition were especially consistent with the vegetation transition across the landscape. Glomeromycetes increased and Agaricomycetes declined from forest to marsh, matching the expected shift from EM- to AM-dominated plant communities. Archaeal patterns likewise indicated that hydrology and associated anaerobic conditions were important filters across the gradient, with Methanomicrobia relatively more abundant in marsh soils. Patchy archaeal enrichment outside the marsh further suggests that anaerobic microsites occurred across the landscape, especially in the ecotone where local heterogeneity in water status and organic matter likely generated mixed oxic and anoxic conditions. Overall, taxonomic composition was highly informative for showing how microbial communities were reorganized across the transect, but because these compositional shifts were tightly aligned with the same environmental contrasts already represented in the baseline model, they likely overlapped with rather than extended the predictive information provided by environmental variables.

Gene-based trait descriptors highlighted shifts in decomposition potential across the transect more clearly than they improved prediction of respiration. Y-A-S trait categories were only weakly structured across the gradient, whereas grouped CAZyme abundances differentiated forest and marsh soils in ways that matched the vegetation transition. Lignin-degrading potential was higher in forest and ecotone soils, while cellulose- and hemicellulose-degrading potential were higher in marsh soils, consistent with the contrast between woody and sedge-dominated vegetation (Xiong et al., 2023; Huang et al., 2024). More broadly, the weak performance of broad Y-A-S traits and most grouped CAZyme variables likely reflects the difference between functional potential and realized process rates. Metagenomic profiles captured community gene content and potential decomposition capacity, but functional gene abundance does not necessarily translate directly into realized activity or process rates, as recent work has shown for soil enzyme activities and broader gene-to-process relationships in environmental systems (Rocca et al., 2015; Li et al., 2025). Recent metagenomic studies similarly show that CAZyme profiles can identify vegetation-linked differences in substrate composition and decomposition potential even when they do not directly predict realized process rate (Huang et al., 2024; Yu et al., 2024). In our system, these patterns reinforce the distinction between microbial descriptors that track environmental and substrate structuring and those that improve prediction of realized carbon flux.

## Conclusions

Microbial representation in carbon cycle models is essential to improving predictions of soil respiration, and identifying useful microbial predictors requires testing them across different spatial scales and ecosystems. This study shows that a simple biomass-partitioning metric, the F:B ratio, can improve prediction of heterotrophic soil respiration across a heterogeneous landscape gradient. Added biological detail in the form of taxonomic and gene-based functional trait data were informative for revealing how landscape position and environmental filtering reorganized microbial communities across the forest-to-marsh transition, but these variables did not improve prediction of realized carbon flux because their explanatory value largely overlapped with the same hydrologic and substrate gradients already captured by the environmental baseline model. Meanwhile, F:B ratio was only weakly structured by environmental variables along the gradient. Its predictive power likely stemmed from its integration of community characteristics that influence rate limiting steps in decomposition and the fate of microbial C after uptake, including differences between fungal and bacterial enzymatic capabilities and necromass chemistries. Together, these findings highlight the importance of distinguishing between microbial variables that track environmental structure and those that improve prediction of realized carbon flux. The ecotone was especially informative in this regard, because its overlapping environmental filters and high microsite heterogeneity illustrated how strong community turnover can coexist with weak categorical differences in respiration. For future studies, this argues for sampling designs that explicitly span ecotones and that test microbial predictors against environmental baselines rather than treating added biological detail as inherently predictive.

## Acknowledgements

We acknowledge funding from the US National Science Foundation (award no. 2319597). Site access to Kunz Marsh was kindly permitted by the South Slough National Estuarine Research Reserve in Oregon, USA. We thank the following individuals for their assistance with fieldwork and lab work: Emily Huckstead, Caryn Iwanaga, Katya Podkovyroff, Rachel Dennis, and Karen Adair.

## Author Contributions

CEM conceptualized and designed the experiment with input and supervision from LRA, BJMB, JR, and LCRS. CEM performed fieldwork and laboratory analyses, conducted bioinformatic analyses and all statistical analysis, and drafted the original manuscript. LRA, BJMB, JS, and LCRS acquired funding. All authors were involved in critical revision and approval of the final manuscript.

## Conflict of Interest Statement

The authors declare no conflicts of interest.

## Notes

### Competing Interest Statement

The authors have declared no competing interest.

