## Appendix S1 - Tables and Figures for "Fungal:bacterial biomass balance links environmental gradients to soil respiration across a forest-to-marsh transition"

**S1 Table 1:** CAZyme families were assigned to biologically defined origin and substrate groups to support grouped analyses of abundance patterns associated with plant-, fungal-, and bacterial-derived organic matter pools. Table data adopted from López-Mondéjar et al. (2020).

| Biomass origin | Substrate group | CAZy family | Activity |
|---|---|---|---|
| Bacterial | Peptidoglycan | GH102 | peptidoglycan lytic transglycosylase |
| Bacterial | Peptidoglycan | GH103 | peptidoglycan lytic transglycosylase |
| Bacterial | Peptidoglycan | GH104 | peptidoglycan lytic transglycosylase |
| Bacterial | Peptidoglycan | GH108 | lysozyme |
| Bacterial | Peptidoglycan | GH22 | lysozyme |
| Bacterial | Peptidoglycan | GH23 | lysozyme/peptidoglycan lytic transglycosylase |
| Bacterial | Peptidoglycan | GH24 | lysozyme |
| Bacterial | Peptidoglycan | GH25 | lysozyme |
| Bacterial | Peptidoglycan | GH73 | peptidoglycan hydrolase with endo-beta-N-acetylglucosaminidase specificity |
| Fungal | Chitin | AA11 | lytic polysaccharide monooxygenase |
| Fungal | Chitin | GH18 | chitinase |
| Fungal | Chitin | GH19 | chitinase |
| Fungal | Chitin | GH20 | N-acetyl beta-glucosaminidase |
| Fungal | Glucans | GH128 | endo-1,3-beta-glucanase |
| Fungal | Glucans | GH17 | endo-1,3-beta-glucanase |
| Fungal | Glucans | GH55 | exo-beta-1,3-glucanase/endo-1,3-beta-glucanase |
| Fungal | Glucans | GH64 | endo-1,3-beta-glucanase |
| Fungal | Glucans | GH81 | endo-1,3-beta-glucanase |
| Plant | Cellulose | AA10 | lytic polysaccharide monooxygenase |
| Plant | Cellulose | AA9 | lytic polysaccharide monooxygenase |
| Plant | Cellulose | GH1 | beta-glucosidase |
| Plant | Cellulose | GH116 | beta-glucosidase |
| Plant | Cellulose | GH12 | endoglucanase |
| Plant | Cellulose | GH3 | beta-glucosidase |
| Plant | Cellulose | GH45 | endoglucanase |
| Plant | Cellulose | GH48 | reducing end-acting cellobiohydrolase/endoglucanase |
| Plant | Cellulose | GH5 | beta-glucosidase/endoglucanase |
| Plant | Cellulose | GH6 | cellobiohydrolase |
| Plant | Cellulose | GH7 | reducing end-acting cellobiohydrolase |

| | | | |
|---|---|---|---|
| Plant | Cellulose | GH8 | endoglucanase/endoxylanase |
| Plant | Cellulose | GH9 | endoglucanase |
| Plant | Hemicellulose | CE1 | acetyl xylan esterase |
| Plant | Hemicellulose | CE12 | acetyl xylan esterase |
| Plant | Hemicellulose | CE15 | acetyl xylan esterase |
| Plant | Hemicellulose | CE16 | acetyl xylan esterase |
| Plant | Hemicellulose | CE2 | acetyl xylan esterase |
| Plant | Hemicellulose | CE3 | acetyl xylan esterase |
| Plant | Hemicellulose | CE4 | acetyl xylan esterase |
| Plant | Hemicellulose | CE5 | acetyl xylan esterase |
| Plant | Hemicellulose | CE6 | acetyl xylan esterase |
| Plant | Hemicellulose | CE7 | acetyl xylan esterase |
| Plant | Hemicellulose | GH10 | endoxylanase |
| Plant | Hemicellulose | GH11 | endoxylanase |
| Plant | Hemicellulose | GH115 | xylan alpha-1,2-glucuronidase |
| Plant | Hemicellulose | GH120 | beta-xylosidase |
| Plant | Hemicellulose | GH131 | exo-beta-1,3/1,6-glucanase/endo-beta-1,4-glucanase |
| Plant | Hemicellulose | GH16 | xyloglucanase/endoglucanase |
| Plant | Hemicellulose | GH2 | beta-galactosidase/beta-glucuronidase |
| Plant | Hemicellulose | GH26 | endomannanase |
| Plant | Hemicellulose | GH30 | endoxylanase/beta-1,6-glucanase/beta-xylosidase |
| Plant | Hemicellulose | GH36 | alpha-galactosidase |
| Plant | Hemicellulose | GH39 | beta-xylosidase/alpha-L-arabinofuranosidase |
| Plant | Hemicellulose | GH43 | beta-xylosidase/endoxylanase |
| Plant | Hemicellulose | GH44 | xyloglucanase/endoglucanase |
| Plant | Hemicellulose | GH51 | alpha-L-arabinofuranosidase |
| Plant | Hemicellulose | GH52 | beta-xylosidase |
| Plant | Hemicellulose | GH54 | alpha-L-arabinofuranosidase |
| Plant | Hemicellulose | GH62 | alpha-L-arabinofuranosidase |
| Plant | Hemicellulose | GH67 | xylan alpha-1,2-glucuronidase |
| Plant | Hemicellulose | GH74 | xyloglucanase |
| Plant | Hemicellulose | GH95 | alpha-L-fucosidase/alpha-L-galactosidase |
| Plant | Lignin | AA1 | laccase |
| Plant | Lignin | AA2 | peroxidase |
| Plant | Lignin | AA3 | oxidase |
| Plant | Lignin | AA4 | oxidase |
| Plant | Lignin | AA5 | oxidase |
| Plant | Lignin | AA6 | 1,4-benzoquinone reductase |

**S1 Table Notes:** Biomass origin denotes the broad biomass source with which each CAZyme family was associated, and substrate group denotes the primary substrate category targeted by that family in downstream grouped abundance analyses.

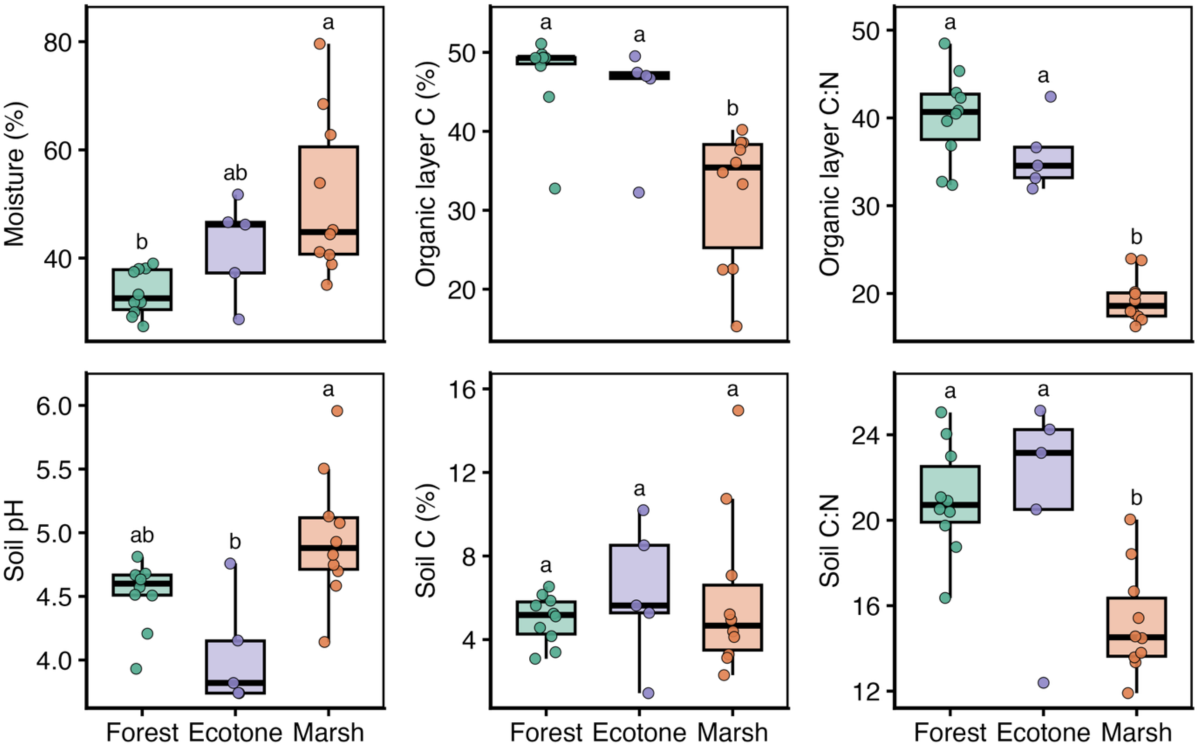


**S1 Figure 1:** Environmental variation across the forest-to-marsh landscape gradient. Boxplots display raw values for soil moisture, organic layer C, organic layer C:N, soil pH, soil C, and soil C:N across landscape positions. Letters indicate Tukey HSD significance groups from ANOVA performed on transformed values.

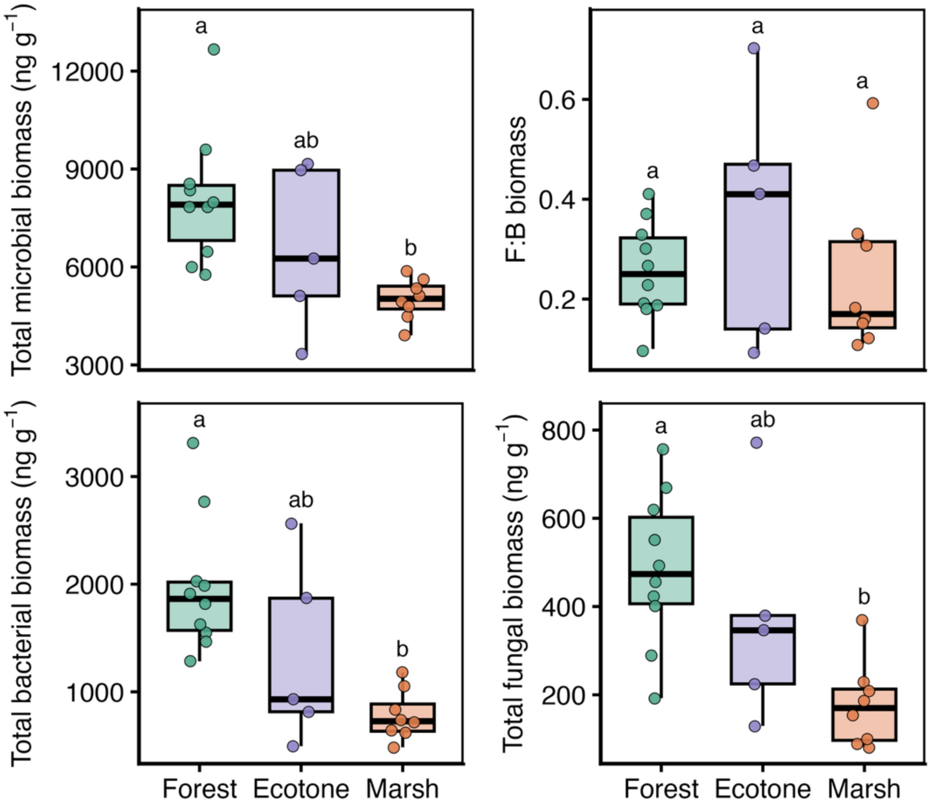


**S1 Figure 2:** PLFA biomass variation across the forest-to-marsh landscape gradient. Boxplots display raw values for total microbial biomass, F:B biomass ratio, total fungal biomass, and total bacterial biomass across landscape positions. Letters indicate Tukey HSD significance groups from ANOVA performed on transformed values.

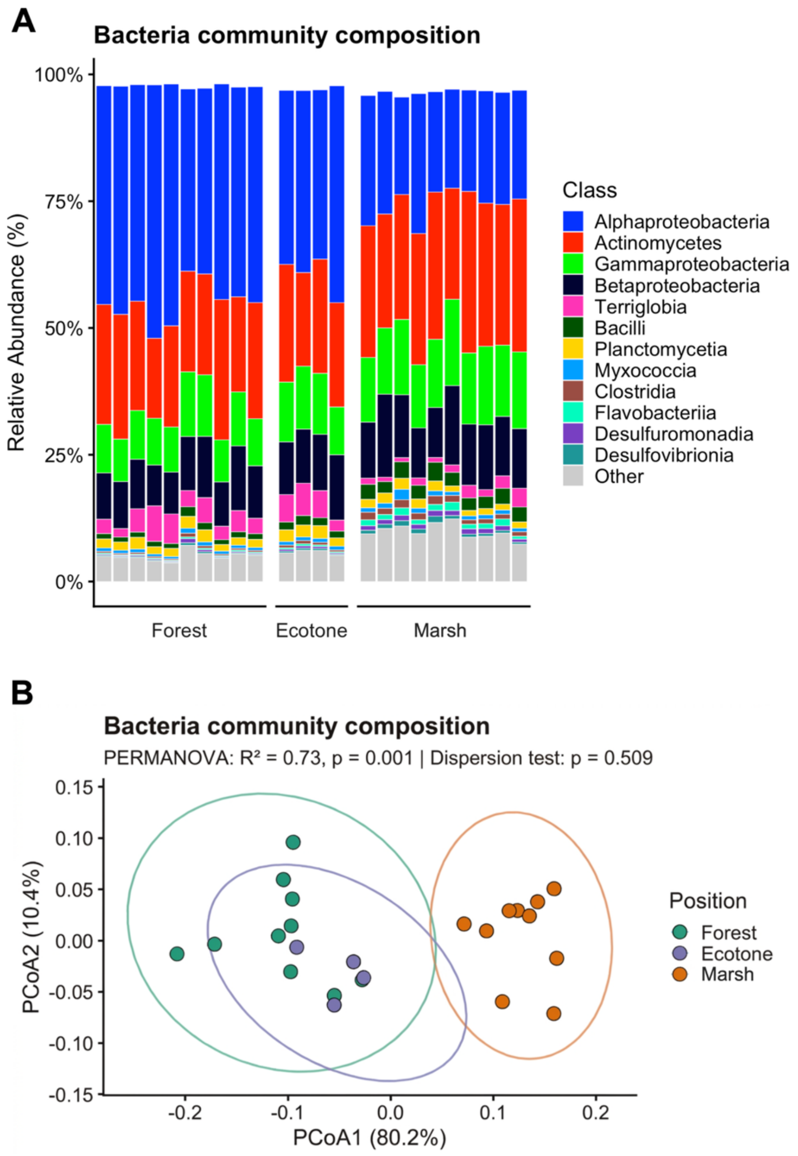


**S1 Figure 3**: Bacterial community composition across landscape positions. (A) Relative read abundance of the 12 most abundant bacterial classes in each sample, grouped by landscape position; all remaining classes are combined as Other. (B) Bray-Curtis ordination (PCoA) of bacterial community composition across landscape positions. Ellipses show the 95% confidence regions for each position. PERMANOVA indicated a significant effect of landscape position on bacterial community composition ($R^2 = 0.73$, $p = 0.001$), while multivariate dispersion did not differ significantly among positions ($p = 0.509$).

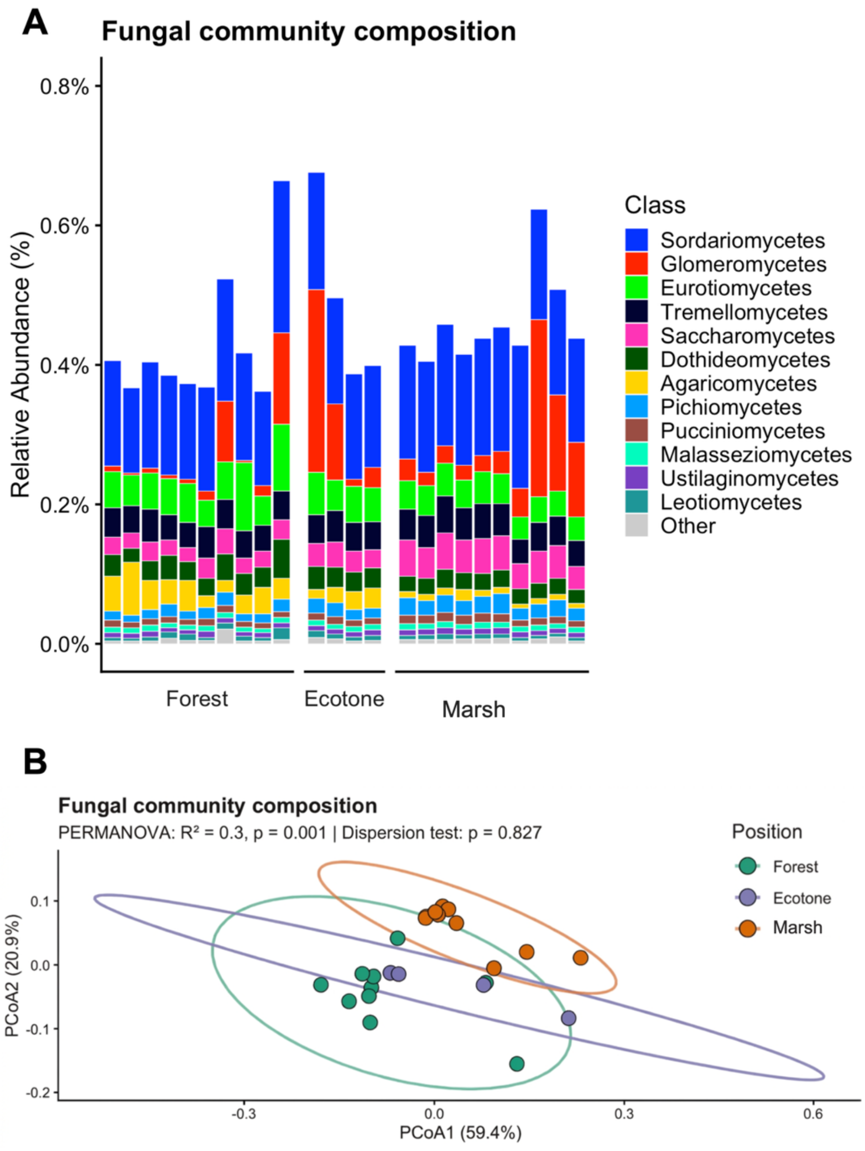


**S1 Figure 4:** Fungal community composition across landscape positions. (A) Relative read abundance of the 12 most abundant fungal classes in each sample, grouped by landscape position; all remaining classes are combined as Other. (B) Bray-Curtis ordination (PCoA) of fungal community composition across landscape positions. Ellipses show the 95% confidence regions for each position. PERMANOVA indicated a significant effect of landscape position on fungal community composition ($R^2 = 0.30$, $p = 0.001$), while multivariate dispersion did not differ significantly among positions ($p = 0.827$).

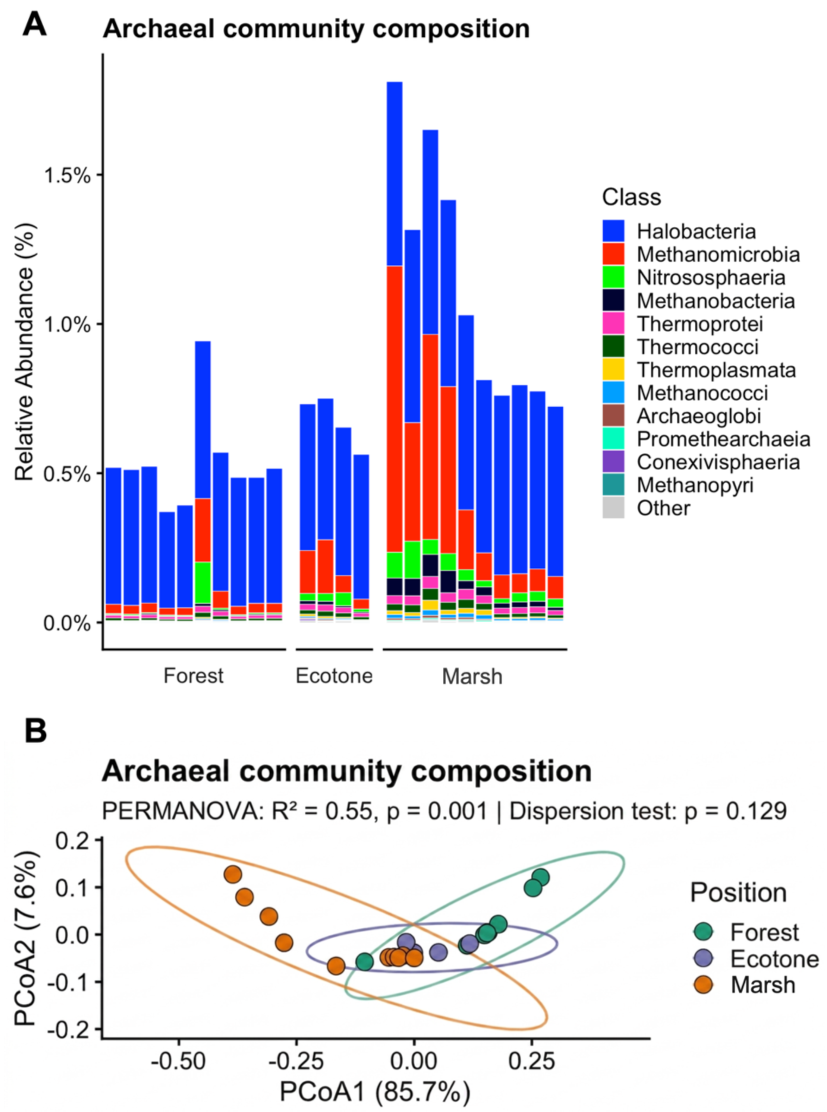


**S1 Figure 5:** Archaeal community composition across landscape positions. (A) Relative read abundance of the 12 most abundant archaeal classes in each sample, grouped by landscape position; all remaining classes are combined as Other. b) Bray-Curtis ordination (PCoA) of archaeal community composition across landscape positions. Ellipses show the 95% confidence regions for each position. PERMANOVA indicated a significant effect of landscape position on archaeal community composition ($R^2 = 0.55$, $p = 0.001$), while multivariate dispersion did not differ significantly among positions ($p = 0.129$).

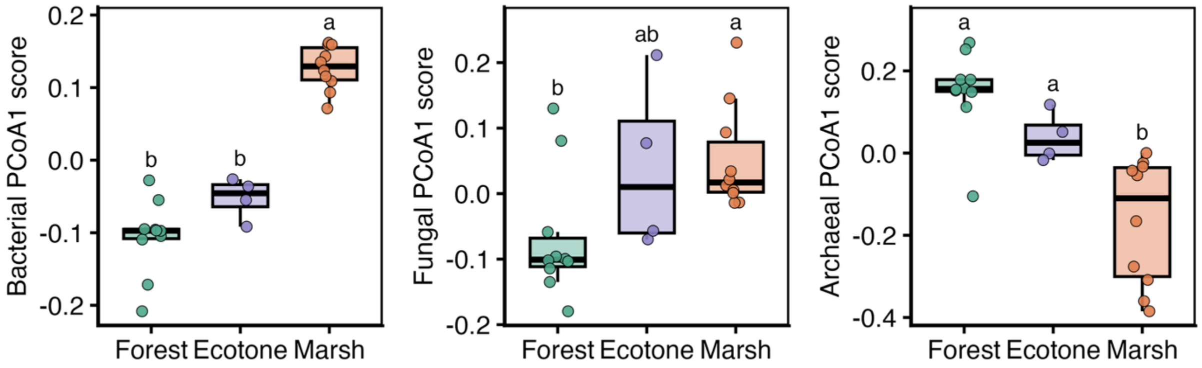


**S1 Figure 6:** Microbial community ordination scores across landscape positions. Boxplots show PCoA1 scores for bacterial, fungal, and archaeal community composition across landscape positions. Letters indicate Tukey HSD significance groups from ANOVA.

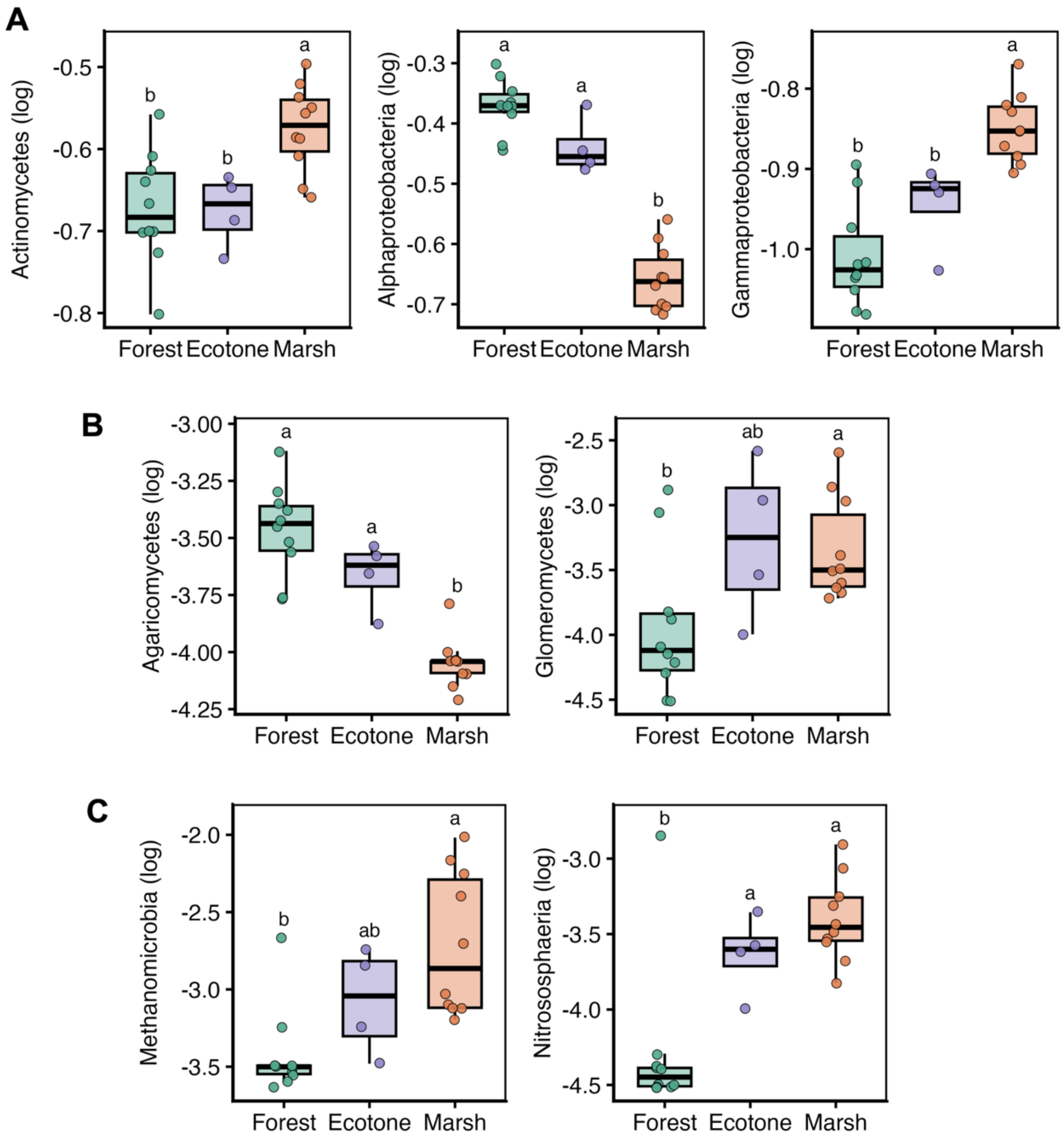


**S1 Figure 7:** Variation in selected taxonomic classes across landscape positions. Boxplots show log-transformed relative abundances of selected (A) bacterial, (B) fungal, and (C) archaeal classes across landscape positions. Letters indicate Tukey HSD significance groups based on ANOVA.

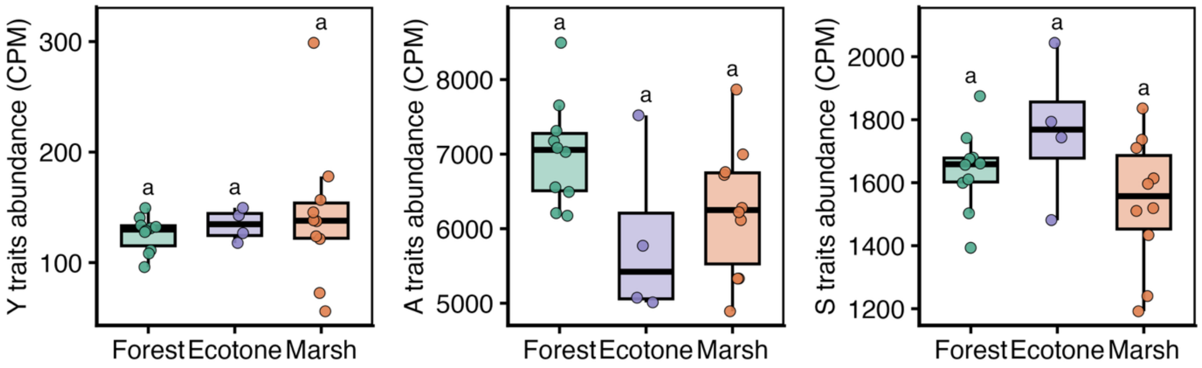


**S1 Figure 8:** Y-A-S trait abundance across landscape positions. Boxplots show raw counts per million (CPM) for Yield (Y), Resource acquisition (A), and Stress tolerance (S) trait categories across landscape positions. Letters indicate Tukey HSD significance groups based on ANOVA conducted on log-transformed CPM values.

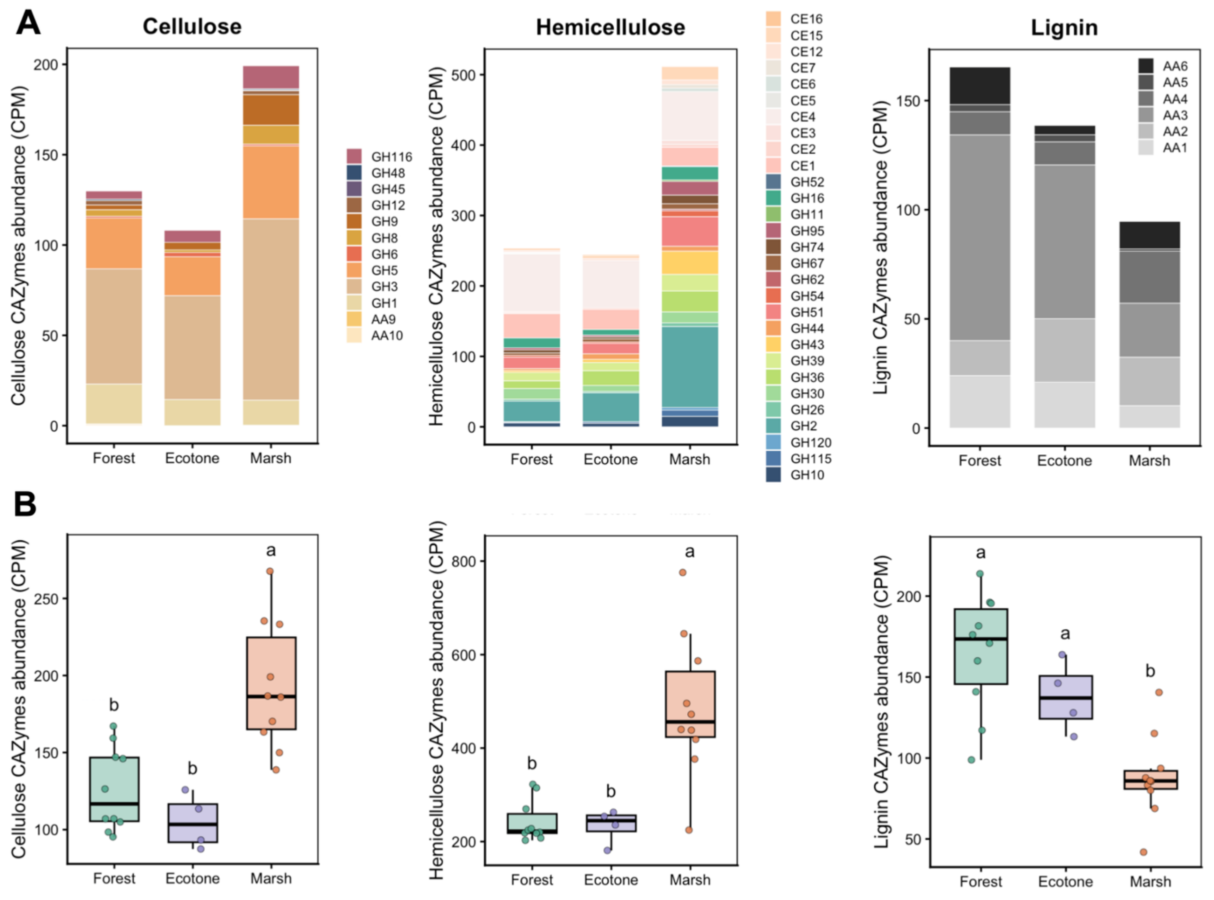


**S1 Figure 9.** Grouped CAZyme abundances associated with degradation of plant biomass across landscape positions. (A) Stacked bars show the mean abundance and family composition of individual CAZyme families, measured as counts per million (CPM), within the cellulose, hemicellulose, and lignin substrate groups across landscape positions. (B) Boxplots show distributions of summed substrate-group abundance as CPM. Letters indicate Tukey HSD significance groups based on ANOVA conducted on log-transformed CPM values.

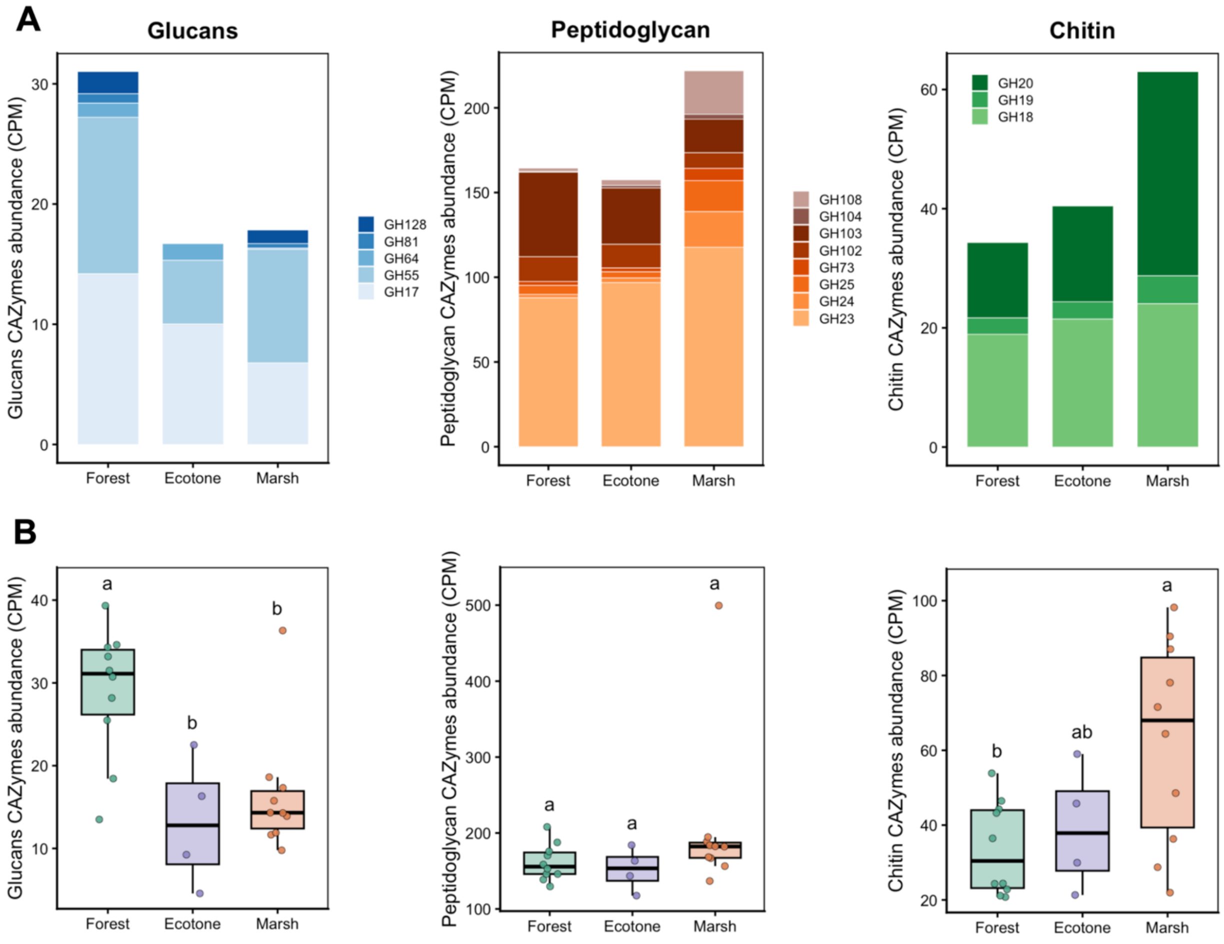


**S1 Figure 10.** Grouped CAZyme abundances associated with fungal- and bacterial-derived biomass across forest, ecotone, and marsh soils. (A) Stacked bars show the mean abundance and family composition of individual CAZyme families, measured as counts per million (CPM), within the glucan, peptidoglycan, and chitin substrate groups across landscape positions. (B) Boxplots show distributions of summed substrate-group abundance as CPM. Letters indicate Tukey HSD significance groups based on ANOVA conducted on log-transformed CPM values.

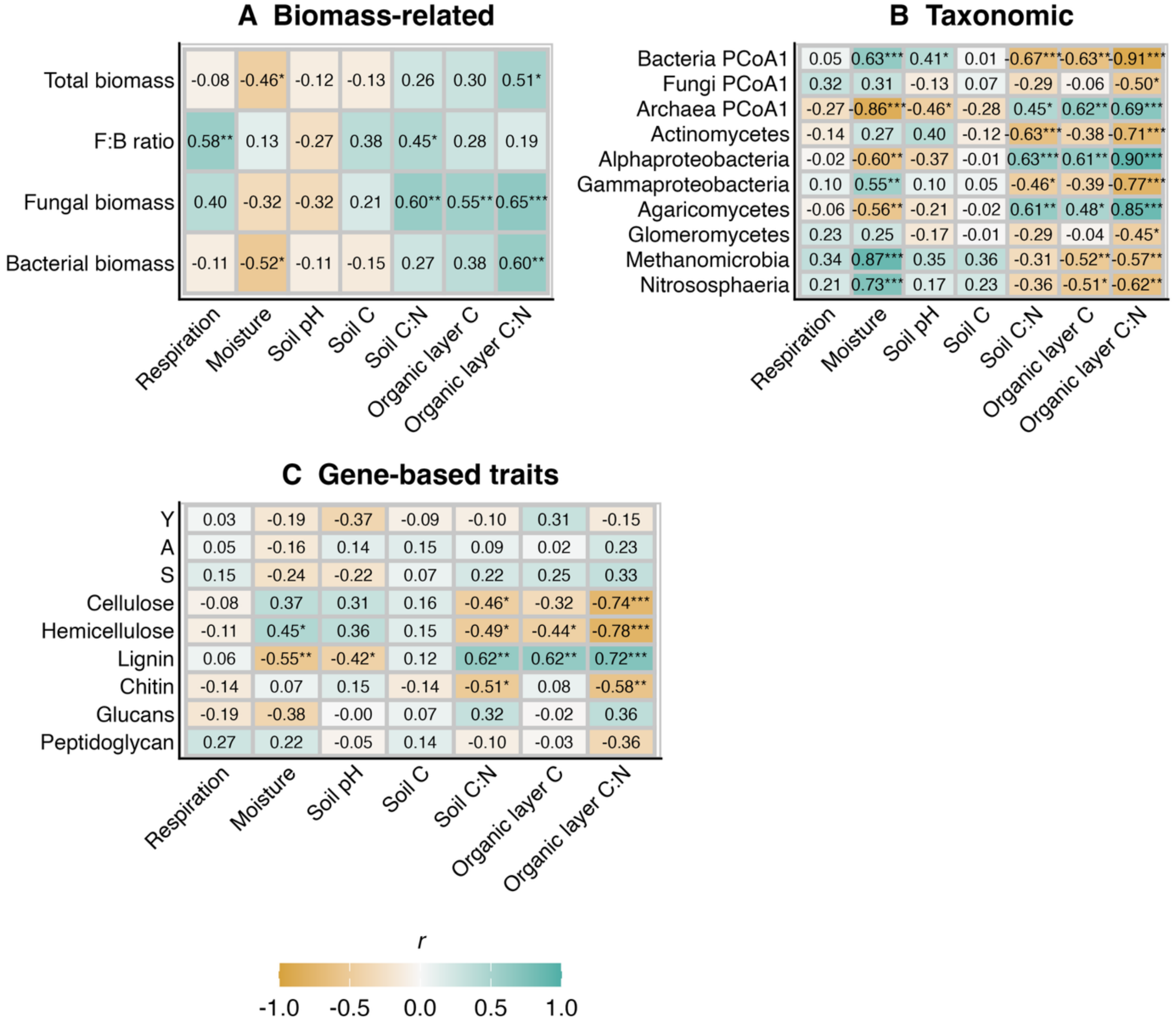


**S1 Figure 11:** Correlation matrices showing associations between environmental variables and (A) biomass-related, (B) taxonomic, (C) and gene-based trait microbial characteristics. Values in each cell are Pearson correlation coefficients (*r*) calculated using log-transformed soil heterotrophic respiration (original units: µmol $CO_2$ $g^{-1}$ $hr^{-1}$) , environmental predictors, and microbial variables. Asterisks indicate significance (* $p < 0.05$, ** $p < 0.01$, *** $p < 0.001$).

**S1 Table 2:** Changes in model support and explanatory power after adding individual microbial predictors to the primary environmental baseline model (soil moisture + organic layer C) explaining soil respiration.

| Response group | Microbial predictor | n | full $R^2_{adj}$ | delta $R^2_{adj}$ | delta AICc | p value |
|---|---|---|---|---|---|---|
| Biomass-related | F:B ratio | 23 | 0.692 | 0.057 | -1.764 | 0.0433 |
| Taxonomic | Methanomicrobia | 24 | 0.687 | 0.037 | -0.601 | 0.0776 |
| Biomass-related | Fungal biomass | 23 | 0.664 | 0.03 | 0.185 | 0.113 |
| Taxonomic | Archaea PCoA1 | 24 | 0.664 | 0.013 | 1.138 | 0.192 |
| Taxonomic | Nitrososphaeria | 24 | 0.658 | 0.008 | 1.533 | 0.24 |
| Gene-based traits | Hemicellulose | 24 | 0.658 | 0.007 | 1.562 | 0.244 |
| Gene-based traits | Chitin | 24 | 0.657 | 0.007 | 1.595 | 0.249 |
| Taxonomic | Fungi PCoA1 | 24 | 0.654 | 0.003 | 1.832 | 0.287 |
| Gene-based traits | Peptidoglycan | 24 | 0.652 | 0.002 | 1.953 | 0.309 |
| Gene-based traits | S traits | 24 | 0.651 | 0 | 2.049 | 0.328 |
| Gene-based traits | A traits | 24 | 0.648 | -0.002 | 2.212 | 0.364 |
| Gene-based traits | Cellulose | 24 | 0.648 | -0.003 | 2.251 | 0.373 |
| Taxonomic | Glomeromycetes | 24 | 0.644 | -0.007 | 2.519 | 0.448 |
| Gene-based traits | Glucans | 24 | 0.639 | -0.012 | 2.84 | 0.574 |
| Taxonomic | Gammaproteobacteria | 24 | 0.637 | -0.014 | 2.977 | 0.652 |
| Gene-based traits | Lignin | 24 | 0.637 | -0.014 | 2.988 | 0.658 |
| Taxonomic | Agaricomycetes | 24 | 0.636 | -0.014 | 3.005 | 0.67 |
| Gene-based traits | Y traits | 24 | 0.635 | -0.016 | 3.121 | 0.768 |
| Biomass-related | Total microbial biomass | 23 | 0.618 | -0.017 | 3.194 | 0.763 |
| Taxonomic | Alphaproteobacteria | 24 | 0.633 | -0.017 | 3.214 | 0.915 |
| Taxonomic | Actinomycetes | 24 | 0.633 | -0.017 | 3.216 | 0.921 |
| Taxonomic | Bacteria PCoA1 | 24 | 0.633 | -0.017 | 3.227 | 0.979 |
| Biomass-related | Bacterial biomass | 23 | 0.617 | -0.018 | 3.234 | 0.808 |
